# Characterization and implication of the Orai3 channel and ABC type transporters in the phenomenon of chemoresistance to cisplatin and pemetrexed in lung cancer

**DOI:** 10.1101/2024.09.22.613742

**Authors:** Daoudi Redoane

## Abstract

Orai3 channels have been associated with cell proliferation, survival and metastasis in several cancers. Previous studies have shown that Orai3 seems to be involved in the development of acquired chemo-resistance of cisplatin in non-small cell lung cancer via ABC transporters that transport certain chemotherapeutic agents, such as cisplatin, out of the cells. Based on these studies, we hypothesize that Orai3 may be involved in the development of acquired chemo-resistance of pemetrexed. By using MTT assay, we show that pemetrexed is efficient (IC50 =0,4*µM*)and significantly decreases cell viability in a dose dependent manner. Furthermore, we demonstrate using siRNA-mediated Orai3 knockdown that Orai3 silencing has no effect on the chemoresistance against pemetrexed. Then, calcium imaging reveals that pemetrexed doesn’t affect the activity of calcium channels like Orai3. As transcription factors are central to the regulation of gene expression, we wonder if pemetrexed could impact on Orai3 expression. To address the issue,*Mn*2+-quench assays are performed in siOrai3 transfected A549 cells incubated with1*µM*pemetrexed for 72h. Finally, RT-PCR is used in order to determine the mRNA expression profile for Orai3 and ABC transporters in A549 cells. We demonstrate that pemetrexed increases Orai3 expression. Taken together, these results suggest that Orai3 is not involved in the development of acquired chemo-resistance of pemetrexed in non-small cell lung cancer. Pemetrexed upregulates Orai3 expression but doesn’t affect the activity of Orai3 channels. As Orai3 channels are known to be involved in the development of acquired chemo-resistance of cisplatin in non-small cell lung cancer, pemetrexed could reduce the therapeutic efficacy of cisplatin if Orai3 protein levels are linked to Orai3 mRNA levels. Moreover, cisplatin has been shown to both increase expression and function of Orai3, resulting in increased chemoresistance. Therefore, our results show that Orai3 could constitute a robust therapeutic target in non-small cells lung cancer.

## INTRODUCTION

Lung cancer is responsible for 1.3 million deaths each year worldwide and remains associated with high mortality in both men and women. To date, clinical studies show that unfortunately the 5-year survival rate does not exceed 5%. The most common clinical signs include difficulty breathing, shortness of breath, weight loss and coughing, possibly accompanied by spitting blood. The most common cause of lung cancer is prolonged exposure to tobacco smoke, including passive smoking.

Forms of cancer are also found in non-smokers and are attributable tothe combination of various factors such as genetics, pollution or agents such as asbestos.

85% of lung cancers are so-called non-small cell cancers. These are subdivided into three subgroups: pulmonary adenocarcinoma (40% of cases), squamous cell carcinoma (40% of cases)and large cell carcinoma (20% of cases). Lung adenocarcinomas are at the center of therapeutic concerns as their incidence continues to increase. The remaining 15% are small cell cancers.

Current treatments differ depending on the type of lung cancer and include radiotherapy, surgeryand chemotherapy. For pulmonary adenocarcinomas, the preferred therapeutic axis is based on the combination of pemetrexed (Alimta )and cisplatin. Cisplatin, a platinum salt, binds to purine bases in DNA cells, altering DNA conformation and thereby inhibiting replication and transcription phenomena,which results in the cessation of cell proliferation. Pemetrexed is a folic acid analogue of the antimetabolite class. Its antiproliferative action is mediated by the inhibition of folate-dependent enzymatic pathways important for replication.

Thus, the enzyme thymidylate synthetase, responsible for the biosynthesis of thymine, is inhibited. Pemetrexed then synchronizes cells in S phase and blocks their proliferation. It has been suggested that pemetrexed may also induce cell death by permanent activation of the AKT pathway, suggesting an atypical proapoptotic role of AKT (*CHEN et al., 2014*).

However, several clinical studies have highlighted phenomena of chemoresistance with respect tothe proposed treatments. Indeed, once in the cells, the active ingredient can be rejected into the extracellular environment, reducing the intracellular concentrations of the drug and therefore its effectiveness. The treatment becomes less effective over time. Chemoresistance results from complex modifications of cellular functioning. Among the different molecular actors studied, numerous clinical studies have highlighted the role of the ABC-type protein family, for example in the efflux of cisplatin into the extracellular environment (*GUMINSKI et al.,2006)*.

This family is composed of two main subgroups: P-glycoproteins and MRPs. P- glycoproteinslocated in the plasma membrane (*JULIANO et al., 2014*) are encoded by genes of the MDR subfamily. These proteins are expressed in a large number of cell types such as enterocytes, endothelial cells of the blood-brain barrier or even certain cancer cells. They confer increased resistance of cells to cytotoxic molecules as well as to many drugs (*AMBUDKAR et al., 1999*).

MRPs, like P-glycoproteins, are ATPase transporters (*COLE et al., 1992*).

It has been shown that loss of MRP1 leads to a return of sensitivity of lung cancer cells to chemotherapies. In addition, expressed in NSCLC cells, MRP leads to resistance to certain anticancer molecules (*GIACCONE et al., 1996*).MRP-type proteins such as P-glycoproteins thereforehave the capacity to cause an efflux of cisplatin, thereby causing a reduction in the efficacy of the cytotoxic agent.

Many signaling pathways that regulate the expression of these proteins are thus involved in the phenomenon of chemoresistance. Among these, the PI3K pathway plays a major role. The kinaseactivity of PI3K causes the activation of proteins of the AKT family (*FRANKE et al., 1995*).

The MAPK pathway also appears to be involved in chemoresistance processes. Indeed, it appears that the mutation of a gene of the RAS family, KRAS, is found in 15 to 30% of adenocarcinomas. Thismutation confers constitutive activity to the RAS/MAPK pathway (*RODENHUIS et al., 1988*).

Physiologically, activation by a growth factor of a receptor with tyrosine kinase activity allows theauto-phosphorylation of the receptor at tyrosine residues by conformational change and releaseof a site with kinase activity on the cytoplasmic domain of the receptor. The phosphorylated and possibly dimerized receptor then allows the recruitment of the adaptor protein GRB2 via a SH2 domain of GRB2.

GRB2 recruits SOS via its SH3 domain. SOS is a Ras-GEF which allows the activation of Ras via the exchange of Ras-GDP into Ras-GTP. The extinction of the pathway will involve a Ras-GAP which will promote the GTPasic activity of Ras to give back inactive Ras-GDP.Subsequently Ras-GTP causes the activation of Raf indirectly then Raf activates MEK proteins which activate by phosphorylation members of the MAPK family such as Erk1 or Erk2. Once phosphorylated, Erk proteins are translocated into the nucleus where they phosphorylate transcription factors such as c-fos, c-jun or c-myc, playing a primordial role in cell proliferation.

Moreover, it is now well established that Ras can recruit and activate PI3K at the membrane, showing atrue functional link between the MAPK and PI3K/AKT pathways. AKT being involved by the continued in ABC-dependent chemoresistance since it has been shown that AKT increases the level of primary transcripts and the functionality of certain ABC transporters involved in chemoresistance phenomena. As such, a mutation of the KRAS gene is correlated with an increasein cell proliferation but also with increased resistance of cancer cells to cytotoxic treatments.

It is necessary to distinguish between innate chemoresistance, which is present even in the absence of treatment,and acquired chemoresistance, which is promoted and amplified by the administration of treatment.

The MAPK pathway as described above but also on other more specific mechanisms present at certain stages or locations of tumors only, such as hypoxia. Concerning the mechanisms of acquired chemoresistance, recent data suggest a link between cisplatin, the increase in the level of primary transcripts of the Orai3 channel and an increase in the phenomenon of acquired chemoresistance. Orai3 is a SOC-type membrane calcium channel that opens following emptying of calcium stores in the endoplasmic reticulum. There are also the Orai1 and Orai2 isoforms for the Orai family channels. STIM family proteins, reticulum calcium sensors, detect a decrease in calcium levels in the reticulum, tetramerize and physically couple to the Orai3 channel at the plasma membrane, allowing its opening and calcium entry promoting chemoresistance pathwaysvia activation of the AKT pathway and ABC-type transporters.

In addition, previous work shows that the amount of Orai3 increases after a course of cisplatin- based chemotherapy. This work also shows that activation of Orai3 channels is required to allow the activity of the Ras/MAPK and PI3K/AKT signaling pathways, even in the presence of the KRASmutation (*AY et al., 2013*).This data suggests that in the context of innate chemoresistance a basallevel of Orai3 channel activity is necessary for cancer cell proliferation and the resistance phenotype. In the context of acquired chemoresistance this shows that cisplatin administration contributes to increasing the resistance of cancer cells to cisplatin itself.

All of these data allow us to note that in the processes of tumorigenesis, calcium is a veryimportant second messenger, intervening both in the mechanisms of proliferation and chemoresistance of cancer cells.

With regard to pemetrexed, it is well established that it activates AKT to promote apoptosis, whichat the same time amplifies chemoresistance dependent on ABC-type transporters and mediated through AKT.

On the other hand, to our knowledge, there is no study that has proven an interaction between the pemetrexed and the Orai3 channel, which is already known to be involved in chemoresistance tocisplatin.

Based on the previous data already established, we assumed that pemetrexed could interact withthe Orai3 channel, in the same way as cisplatin, and therefore be involved in the phenomenon of chemoresistance either towards cisplatin alone or towards cisplatin and pemetrexed itself.

The work that was carried out during this internship had the main objective of highlighting or not a linkbetween pemetrexed and the Orai3 channel and of specifying the nature of the interactions between pemetrexed and the Orai3 channel if such a link existed.

To conduct this entire study, the A549 cell line was used. These are human alveolar

Epithelial cancer cells (non-small cell adenocarcinoma).

## MATERIALS AND METHODS

### Culture experiments

Cell culture experiments were performed in a category 2 culture room,under a type II biological safety hood, BH 2004 D, with vertical laminar flow.

### Preparation of cell culture medium

The culture medium used was prepared on the basis of a volumeof 500 mL and was composed of: 385.4 mL of sterile water, 50 mL of EMEM 10X, 14.6 mL of bicarbonates 7.5%, 10 mL of HEPES 1M, 5 mL of non-essential amino acids (MEM 100X), 5 mL of gentamicin, 5 mL of L-glutamine 200 mM, 25 mL of fetal calf serum.

### Preparation of cells

The A549 cell line was used. These are human alveolar epithelial cancer cells (adenocarcinoma). These cells have a mutation in the KRAS gene, which makes the Ras protein constitutively hyperactive due to a lack of GTP hydrolysis into GDP with exaggerated activation ofthe MAPK and PI3K/AKT pathways. This cell type was chosen for our study because it representscases where the chances of chemotherapy treatment being effective are the lowest.

Ampoules containing A549 cells were maintained in liquid nitrogen (-80°C) and then thawed in awater bath at 37°C. The cells were then suspended in 9 mL of culture medium and centrifuged (1000 rpm for 10 minutes).

After centrifugation and to remove the freezing medium, the supernatant obtained was removedand the A549 cells were resuspended in 1mL of culture medium.

The suspension was placed in a 25cm flask containing 5mL of culture medium. This flask was thenkept in an incubator containing 5%*CO*2, an atmosphere saturated with humidity and at 37°C. After 24 hours of incubation, the cells adhere to the bottom of the flask and proliferate, forming a cell carpet. The medium was then renewed every 48 hours to optimize cellproliferation.

After 6 to 7 days of culture, the cells occupy 80% of the surface of the flask, justifying detachment of the cells by means of trypsin treatment to avoid inhibition of proliferation by contact and thusavoid distorting the results of subsequent experiments.

### Detachment of cells

The culture mediumwas removed using an aspiration syringe and then two successive washes with PBS were carried out in order to eliminate the trypsin inhibitors present in the serum.

800*µL*Trypsin was then applied to the cells, together with a divalent cation chelator, EDTA. Thisstep allowed the cells to be detached from the culture dish.

After a few minutes of waiting, the previously applied trypsin was neutralized by the addition of 10mL of culture medium containing 10% fetal calf serum, refluxes were carried out to homogenizethe cell suspension.

### Counting on Malassez cell

In order to determine the volume of cells to be deposited in each wellor dish for the different tests used in this study (MTT, calcium imaging, real-time quantitative RT- PCR), a cell count was carried out on Malassez cells.

### Transfections

The culture medium was initially removed by aspiration. A PBS rinse was performed to remove the remaining culture medium in the flask.

800*µL*of trypsin were added to detach the cells. 2 1.5mL eppendorfs (1 eppendorf SiCtrl, 1eppendorf SiOrai3) each containing106cells were prepared.

The eppendorfs were then centrifuged (1000 rpm for 7 minutes, 22 C°). The supernatant wasaspirated and the pellet containing the cells was retained.

100*µL*of nucleofactant were added into the eppendorfs to resuspend the pellet and then 2*µg* SiCtrl and SiOrai3 were added, each in a different eppendorf. All the medium contained in the eppendorfs was collected and then deposited in a transfection tank (1 SiCtrl tank, 1 SiOrai3 tank).

Transfection was then performed using the Nucleofector II transfection apparatus (electroporation transfection) set with the transfection program adapted to the A549 cell line.500 *µL* of culture medium was added to each of the two tanks to restore the cells to favorable conditions, the medium present in the two tanks was then removed and then transferred back to two separate Eppendorfs. The two Eppendorfs were kept at 37°C for 5 minutes before transferringthe transfected cells to the dishes intended for the different tests of the study, these dishes were placed in an incubator until their use.

### MTT test

The MTT cell viability assay was used. MTT is metabolized by succinate dehydrogenase inthe mitochondria of living cells to form formazan, which forms a purple precipitate, visible at 550 nm. Reading the absorbance at this wavelength thus provides information on the rate of viable cells between several conditions.

50,000 cells were introduced into each dish intended for the MTT test. The cells were detachedagain by the addition of trypsin because the A549 line adheres easily to the new support.

800*µL*of culture medium containing 0.5 mg/mL MTT was added to each dish. MTT was stored inthe dark and refrigerated. The dishes were incubated for 60 minutes and then the medium containing MTT was removed by aspiration.

800*µL*DMSO was added to recover the formazan formed by the living cells and then the mediumcontaining the formazan was transferred to a 96-well plate before reading the absorbance. at 550nm taking into account the background noise of the plate.

The Nanodrop 2000 spectrophotometer was used for automated reading of absorbances in each well, software attached to the device allowed the results to be presented in an excel file.

### Quench- manganese

The ability of the plasma membrane to allow calcium to pass through can be estimated by the manganese quench technique. Indeed, manganese is a divalent ion with a steric hindrance close to that of calcium and therefore uses the same transporters as calcium. Fura-2 (sigma) is a ratiometric probe derived from calcium chelators (EGTA and BAPTA). The probe is made permeant by adding an acetomethylester (AM) group. It can thus cross the plasma membrane but it is not yet able to bind calcium. The probe is indeed activated by the cleavage of its AM group by intracellular esterases, it is then able to bind to ions capable of fixing themselves to it, such as calcium or manganese. The probe is present in free form and bound to calcium, these two forms respond to excitation at 360 nm then the fluorescence intensity is measured at512 nm. Before manganese fixation, the bleaching phenomenon (loss of natural fluorescence of the probe) causes a progressive decrease in fluorescence intensity. The fixation of manganese on fura-2 causes a decrease in fluorescence intensity during excitation of the probe at 360 nm up to astable level where the fluorescence intensity no longer decreases.

Thus, the fluorescence decay kinetics measured at 512 nm during the excitation phases makes it possible to estimate the capacity of the plasma membrane to allow manganese to pass from the extracellular compartment to the intracellular compartment and, by extrapolation, the permeability tocalcium. In the study, the quench was performed over a period of 3 minutes with manganese perfusionfrom 60 seconds. The fluorescence decay kinetics were estimated by the value of the slopes of the linesrepresenting the fluorescence decay over time. This analysis was performed with Origin software and the fluorescence intensity measurement over time was performed with Metafluor software.

### Calcium imaging: variation in intracellular calcium concentration

Variations in intracellular calcium concentrations were estimated with the fura-2 probe and Metafluor software. The probewas excited at 340nm in calcium-bound form and at 380nm in calcium-bound form.

### Real-time quantitative RT-PCR

Cells were contacted with 1mL TRIZOL allowed the cells to be lysed. The cell lysate was then collected and frozen at -80°C for 24 hours.

After drying the pellet was rehydrated with22*µL*of water. The amount of genetic material was estimated by measuring the absorbance with the Nanodrop spectrophotometer. For this,1*µL* samples were introduced into the spectrophotometer and a concentration of total RNA in the solution as well as the purity of total RNA were obtained. Reverse transcription from2*µg*of totalRNA was then performed.

The aim of this step is to synthesize DNA complementary to the RNA in order to allow the polymerase used in qPCR to bind to its substrate. To solutions containing2*µg*of RNA, or0,2*µg/µL* RNA, have been added10*µL*of reaction mix. For10*µL*, the mix contained1*µL*reverse transcriptase,1 *µL*RNase inhibitors,0,8*µL*of dNTP (100nM),2*µL*primers (Reverse and Forward),2*µL*of RT buffer and3 ,2*µL*of sterile water. Experimental data provided with the reverse transcription kit. The reverse transcription cycle began with 10 minutes at 25°C to prepare the samples. Then, the samples werebrought to 35° C for 120 minutes, which allowed the primers to anneal and the cDNA synthesis.

Finally, the samples were brought to 85°C for 5 minutes to stop the synthesis and separate theprimers.

Quantitative PCR reactions were then performed on the LightCycler Roche in 420-well microplates and using the ABsolut kitTMqPCR SYBR Green Mixes (ABgene) according to the supplier’s instructions. The principle of qPCR is to amplify a target gene and labelit in order to quantify it. To do this, a DNA intercalator, SYBR green, was used, which emits fluorescence in contact with double-stranded DNA, with maximum intensity at 550 nm. This fluorescence was detected and measured by the LightCycler microspectrofluorimeter what allowed a quantification of the number of copies of the cDNA sequence of interest. At the end of each cycle, the fluorescence is therefore measured and it is possible to visualize the exponential increase inreal time of the quantity of amplicons generated. Finally, the number of cycles necessary for the fluorescence of the target gene is detectable is inversely proportional to the initial amount of the messenger RNA of interest.

This number of cycles was calculated and compared with the other genes of interest as well as for the same gene of interest between different conditions. The target sequences were amplified from specific primers and using polymerase activating at a high temperature. For each sample, a reaction medium was prepared containing4,6*µL*reaction mix (SYBER green, Taq polymerase, dNTP, enzyme buffer and MgCl2),22*µL*of cDNA diluted 1/20,0,5*µL* from first forward,0,5*µL*of primer Reverse (both to 10)2*µM*) And1,6*µL*of sterile water.

The reference gene used was actin, in order to verify that the variations in the number of cycles obtained for the other genes of interest were indeed due to a difference in the initial quantity of messenger RNA and not to the presence of Taq polymerase inhibitors between two conditions or to thefact that less genetic material was collected between two conditions.

### Statistics

The results of the different experiments were analyzed using a two-way ANOVA test.

The significance threshold was p < 0.05.

## RESULTS AND DISCUSSION

### Pemetrexed has an effect on cell viability

Firstly, the aim was to assess the effect of pemetrexed on A549 cells after 72 hours of incubation at different concentrations of the molecule. For this purpose, an MTT cell viability test was performed. This first pharmacological approach made it possible to draw an effect/dose curve (figure 1) and to determine the IC50 of pemetrexed as well as the dose allowing for maximum action. The observed effect is the decrease in absorbance at 550nm, reflecting the decrease in theamount of formazan formed by living cells and therefore a lower proportion of living cells.

**Figure 1.**
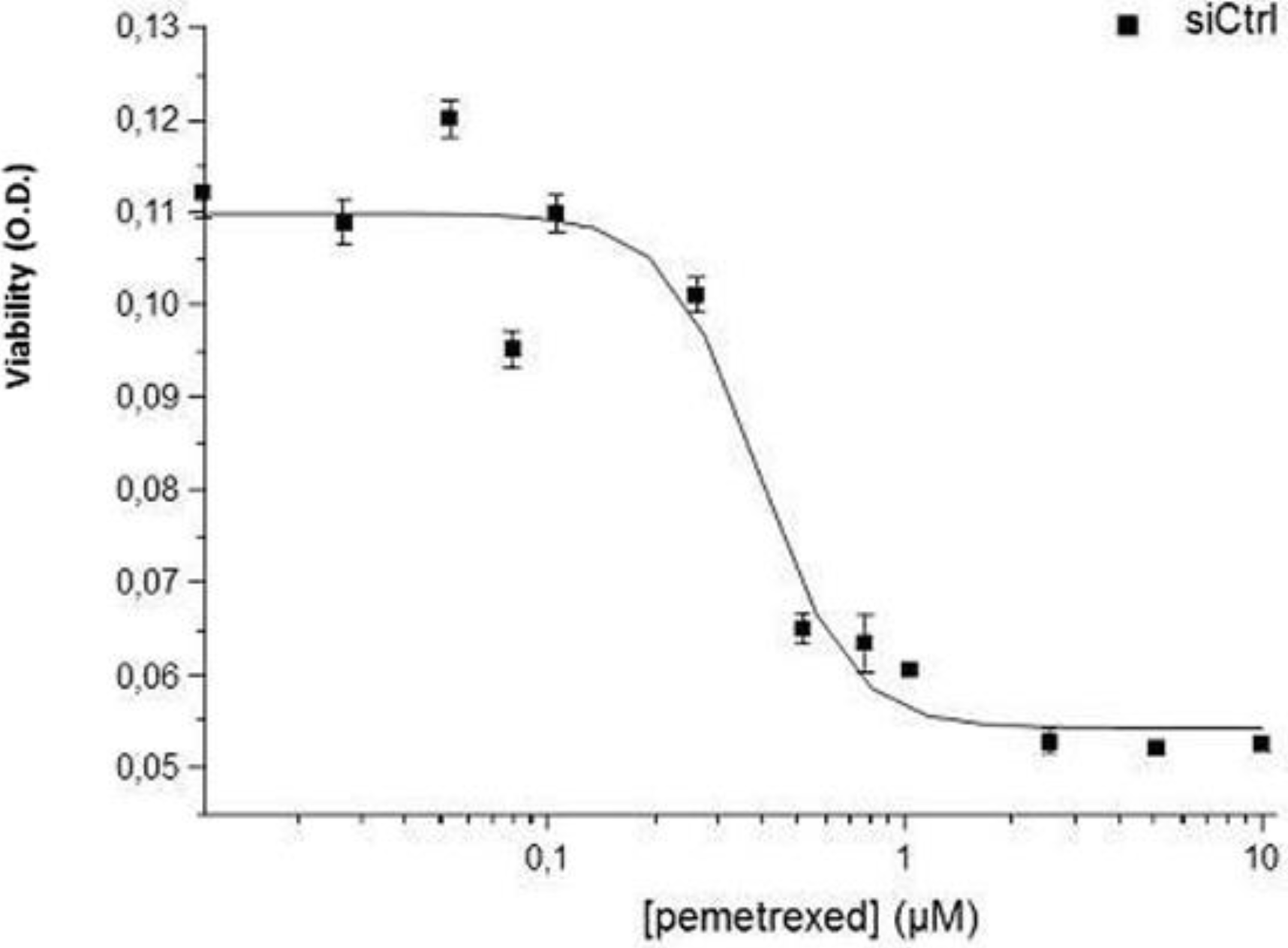
Effect of pemetrexed on cell viability after 72 hours. (n=4)

The decrease in cell viability can be interpreted in three ways. The first is that it can be due to cellmortality, the second to an inhibition of cell proliferation but without mortality, therefore characterized by the same consequence at the absorbance level since fewer cells will be present and therefore less formazan will be formed. Finally, the two phenomena can coexist. As pemetrexed has an effect from a certain dose, it can be assumed that from a certain concentration the mechanisms of resistance to pemetrexed are exceeded, allowing pemetrexed to act. However, the effect of pemetrexed could be observed from a threshold concentration without resistance mechanisms to pemetrexed existing.

The MTT technique is practical and quick to implement. However, it only provides information oncell viability, without specifying the phenomenon causing the decrease in cell viability, i.e. mortality or inhibition of proliferation by synchronization of cells.

Pemetrexed does indeed have an effect on cell viability. We then sought to determine whetherpemetrexed interacted with the Orai3 channel. To do this, we wanted to determine whether pemetrexed was capable of inducing a calcium signal, an important signal in the induction of chemoresistance mechanisms.

### Pemetrexed does not alter calcium channel activity

A first approach therefore consisted of studying the evolution of intracellular calcium homeostasisin the presence of pemetrexed. For this, pemetrexed (1*µM*)was added to the perfusion medium. A first interaction of pemetrexed with calcium channels, therefore possibly with Orai3, was then tested: this involves the modulation of the activity of Orai3 channels already existing at the plasmamembrane. This phenomenon can be observed rapidly in a few minutes if it occurs, unlike regulation of transcription which requires several days. Pemetrexed was thus brought into contact with the cells, i.e. the cells were not previously incubated with pemetrexed. The short-termeffect (15 minutes) of pemetrexed was therefore observed by calcium imaging (figure 2).

**Figure 2.**
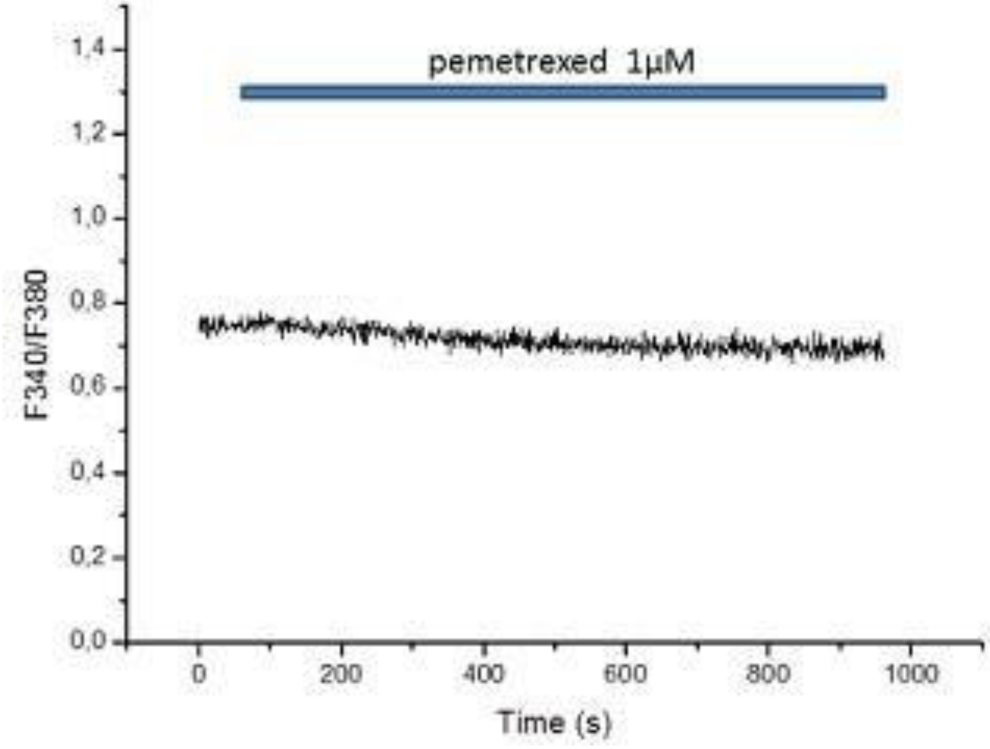
Effect of pemetrexed on calcium channel activity.

The fluoresce intensity after excitation at these wavelengths. Thus there does not appear to be any variation in the intracellular calcium concentration after the application of pemetrexed. If the activity of membrane and/or intracellular calcium channels was modified then the intracellular calcium concentration would have varied, which would have been visible with a variation in the ratio. We therefore wanted to know the effect of pemetrexed on calcium homeostasis after 72 hoursof incubation and thus see if pemetrexed could potentially regulate phenomena other than an increase in the activity of already existing membrane calcium channels.

### Pemetrexed increases membrane calcium permeability

The manganese quench technique was used to visualize the variations in membrane permeability to calcium in the presence and absence of the Orai3 channel, without or with pemetrexed after 72 h of incubation (figure 3).

**Figure 3.**
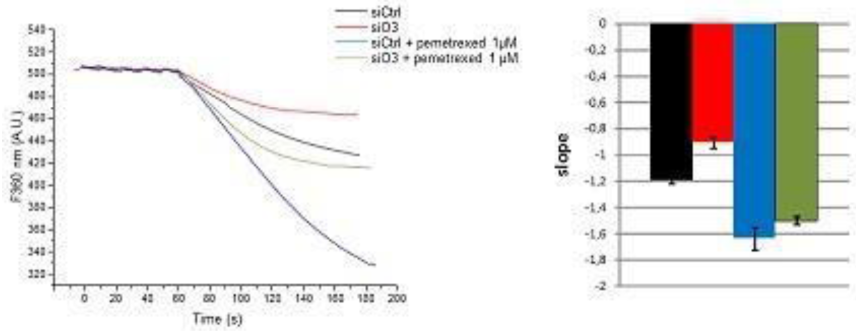
Variations in membrane permeability to calcium with and without Orai3, without or withpemetrexed.

We note that the fluorescence decrease slope is more negative after application of pemetrexed (SiCtrl + pemetrexed compared to the SiCtrl case). This can be explained by the fact that the membrane becomes more permeable to manganese and therefore by extrapolation to calcium in the presence of pemetrexed. This is not due to an increase in calcium channel activity and thereforenecessarily linked to an increase in transcription or membrane addressing (or decreased turnover). To verify the involvement of the Orai3 channel in this phenomenon, functional inhibition of Orai3 using SiOrai3 was performed. Without pemetrexed, the fluorescence decrease slope is more positive for the SiOrai3 case compared to the SiCtrl case, suggesting that the membrane is less permeable to manganese and therefore to calcium due to a smaller number of Orai3 channels. In the presence of pemetrexed and in the case of SiOrai3, membrane permeability to calcium can still be increased. This result suggests that transcription of genes coding for calcium channels other than Orai3 occurs, in addition to that coding for Orai3, and that this is amplified by pemetrexed (1). Pemetrexed could also cause an increase in the expression of Orai3 alone, which cannot be compensated by SiOrai3 (2). A combination of the two phenomena could finally be conceivable (3). In these 3 hypotheses, it is assumed that pemetrexed causes an increase in the transcription of Orai3 (4). Quench is a technique providing information on the functionality of calcium channels since an increase in membrane permeability to calcium is correlated with a greater number of functional membrane calcium channels. Activation of only the already existing channels does not take place. In hypothesis (1) the functional inhibition of Orai3 is supposed to be sufficient to compensate for the increase in transcription of Orai3 by pemetrexed, contrary to hypothesis (2).

However, membrane permeability to calcium is always increased and significantly. If Orai3 does notcontribute to this phenomenon, then it necessarily comes from non-Orai3 calcium channels.

Pemetrexed therefore causes an increase in the transcription of these non-Orai3 calcium channels, but their quantity on the membrane as well as their functionality are also increased; transcription isnot the only parameter amplified. On the other hand, in hypothesis (1), it is impossible to know whether the increase in transcription of Orai3 is associated with a greater number of membrane and functional Orai3s. Indeed, whether or not the increase in Orai3 transcription is linked to a greater quantity of membrane Orai3, membrane permeability to calcium will be increased in the SiCtrl case due to the involvement of non-Orai3 calcium channels.In the case of hypothesis (3) it is also impossible to know whether the increase in the number of functional membrane calcium channels is linked to the increase in transcription. Since in hypothesis (3) this phenomenon can concern only Orai3, only non-Orai3 calcium channels or both types of channels. Only hypothesis (2) allows us to conclude that the increase in transcription of the Orai3 gene is correlated with an increase in the number of functional membrane Orai3.

In fact, in this case it is assumed that pemetrexed increases transcription of the Orai3 gene only. Since membrane permeability is increased in the presence of pemetrexed, this is necessarily due to an increase in the number of membrane and functional Orai3 channels.

It should be noted that the manganese quench technique assumes that manganese can transit via the calcium channels expressed by cells, including Orai3, due to a close steric hindrance for calcium and manganese. However, this remains to be formally demonstrated. Quenchmanganeseis also a rapid technique to implement and provides information on cellular events in real time.

We subsequently wished to verify the involvement of the Orai3 channel in resistance to pemetrexed.

### Resistance to pemetrexed appears to be independent of the Orai3 channel

Functional inhibition of the Orai3 channel by the interfering RNA strategy (SiOrai3) allowed to verify whether the sensitivity of cells to pemetrexed was increased or not in the absence of the functional Orai3 channel. TheMTT assay was used in the presence of SiOrai3 (Figure 4).

**Figure 4.**
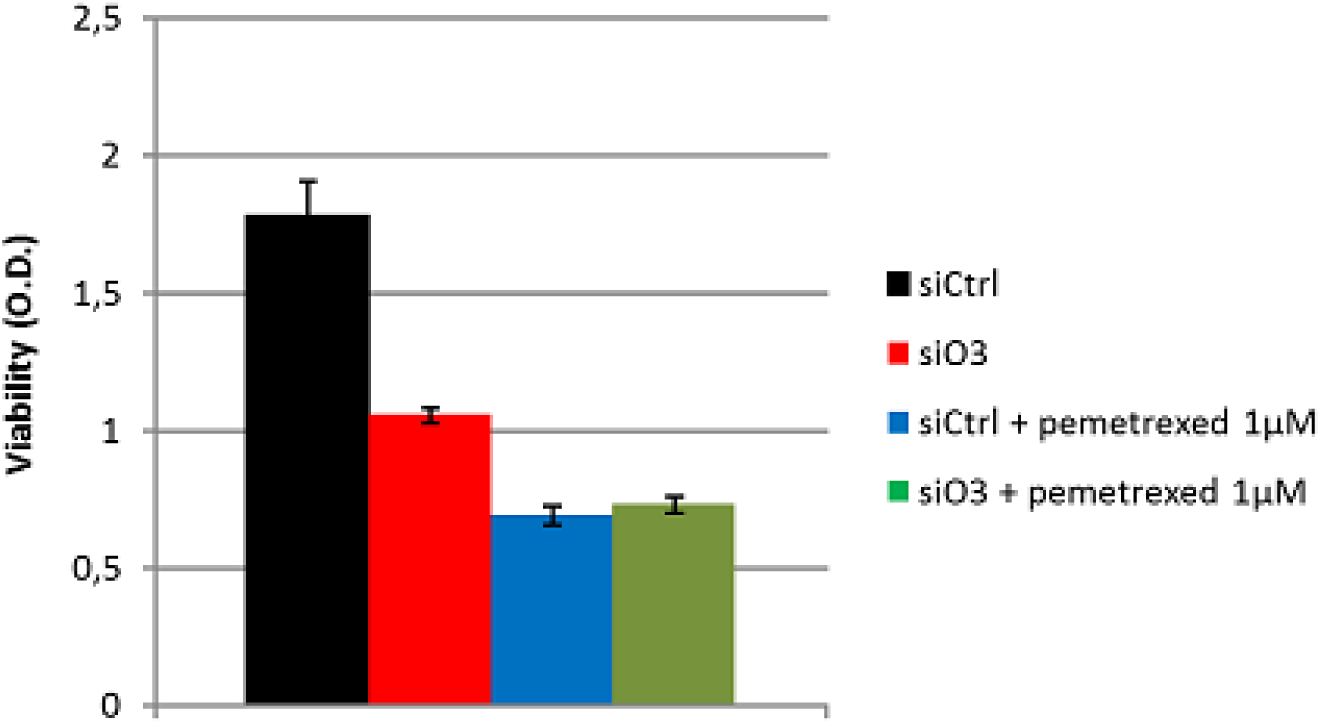
Effect of pemetrexed on cell viability after 72 hours and in the absence of the Orai3 channel.

We observe a decrease in absorbance between the SiCtrl and SiOrai3 cases, which goes from 1.7 to 1. Less formazan is formed in the SiOrai3 case, so fewer cells are viable. This result suggests that Orai3 participates in maintaining the viability of A549, since the functional invalidation of the channel causes a decrease in the latter. This is in accordance with the data already obtained in thelaboratory.

Incubation in pemetrexed1*µM*for 72 hours causes as previously a significant decrease in cell viability. However, we can observe that when Orai3 is invalidated, the same concentration of pemetrexed is required to obtain the same effect on cell viability. This result suggests that the functional invalidation of Orai3 has no effect on sensitivity to pemetrexed. Two hypotheses can beput forward to explain this phenomenon. The number of functional Orai3 channels was much greater at baseline for cells in the SiOrai3 condition compared to cells in the SiCtrl condition and therefore even after functional inhibition of Orai3 the number of Orai3 channels remained identical to that of cells in the SiCtrl condition. In this case, the same concentration of pemetrexedto obtain the same effect would be justified. This first hypothesis can be refuted since the quench highlighted a lower permeability of the membrane to calcium in the SiOrai3 condition, therefore alower number of functional Orai3 channels in the SiOrai3 condition compared to the SiCtrl condition. The second hypothesis is that the number of functional Orai3 channels is indeed reduced in the SiOrai3 case but that other channels potentially involved in resistance to pemetrexed are overexpressed compared to the SiCtrl case, justifying again a greater resistance despite the absence of the Orai3 channel. This hypothesis can be invalidated since the experimentwas repeated 4 times, which is enough to eliminate any sampling fluctuation and variations concerning the number of channels between the two conditions, we thus consider that only the Orai3 channels are reduced in the SiOrai3 case and that the other channels are in equivalent quantity and in the same functionality in the two conditions. There remains only one possible situation. The number of functional Orai3 channels is lower in the SiOrai3 case compared to the SiCtrl case and the other channels involved in pemetrexed resistance are equivalent in number and functionality in both conditions. This implies that if Orai3 were involved in pemetrexed resistance then a lower concentration of pemetrexed would be required in the SiOrai3 case to have the same effect, which is not the case here. We then assume that pemetrexed resistance is independent of the Orai3 channel and that other channels or systems, equivalent in number and functionality in both conditions, are involved in pemetrexed resistance, justifying that despite the functional inhibition of Orai3 the same concentration of pemetrexed is required to have the sameeffect. Since the Orai3 channel is linked to the increase in the activity of certain ABC transporters we hypothesize that pemetrexed is not a substrate of these ABC transporters. Other channels could be responsible for pemetrexed resistance by being coupled to the increase in the activity of other ABC transporters than those regulated by the Orai3 channel or to the activity of other independent ABC resistance systems.

If the other channels were linked to the increase in activity of the same ABC transporters as those regulated by the Orai3 channel then the functional inhibition of Orai3 would have been sufficient to increase the sensitivity of the cells in the SiOrai3 case. Furthermore, the systems responsible for pemetrexed resistance could also be independentof signaling pathways involving non-Orai3 channels.

Finally, there may be no resistance mechanisms to pemetrexed, also justifying that despite the functional inhibition of Orai3, the same concentration of pemetrexed is required to have the same effect.

After showing that resistance to pemetrexed was independent of the Orai3 channel, we wanted to know if pemetrexed could cause an increase in resistance to cisplatin and no longer to itself. To dothis, we sought to know if pemetrexed increased the transcription rate of the Orai3 gene, a channel involved in resistance to cisplatin but not to pemetrexed.

### Pemetrexed increases the transcription rate of the Orai3 gene and of the actors involved in resistance to cisplatin

qPCR was performed for pemetrexed-treated cells after 72 h of incubation (Figure 5).

**Figure 5.**
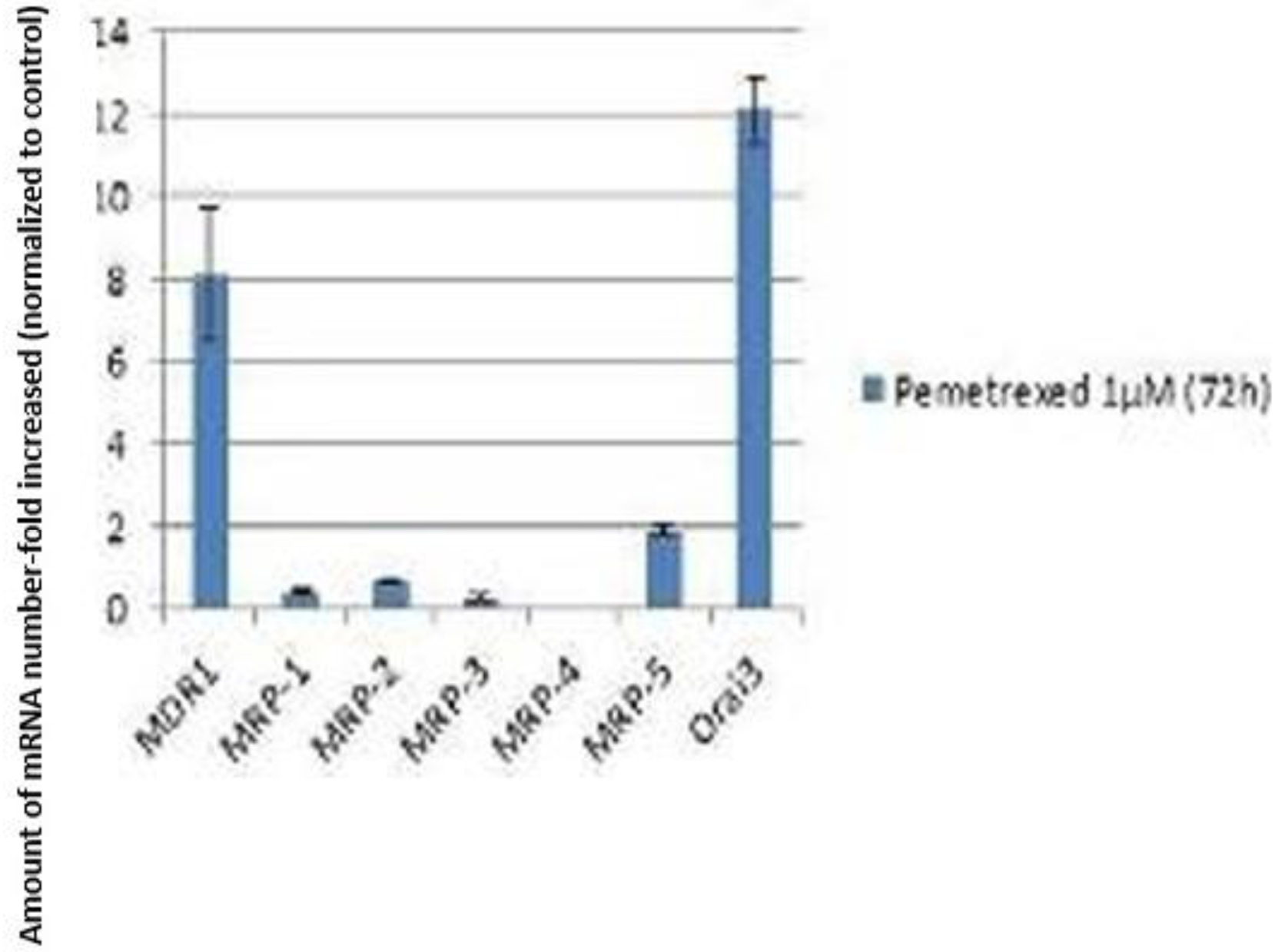
Quantification of the mRNA level of ABC transporters and Orai3 in cells treated with pemetrexed (1µM, 72h)

The amount of Orai3 mRNA is increased 12-fold in the presence of pemetrexed compared to cellsnot treated with pemetrexed. Hypothesis (4) previously formulated during the quench analysis is then validated, pemetrexed increases the transcription rate of the Orai3 gene, without this prejudging an increase in the number of functional Orai3 channels on the membrane. The 3 hypotheses previously formulated during the quench analysis can therefore be retained. The amount of MDR1 mRNA is increased 8-fold in the presence of pemetrexed compared to cells not treated with pemetrexed. It is possible that the increase in the transcription rate of MDR1 is directly linked to that of Orai3 since the calcium transiting through Orai3 allows the increase inthe transcription rate of ABC transporters, including MDR1, via the AKT pathway. It has also been shown that MDR1 is involved in resistance to cisplatin, which could then be amplified by the administration of pemetrexed. Finally, the amount of MRP-5 mRNA is doubled in the presence of pemetrexed compared to cells not treated with pemetrexed. As for MDR1, this result suggests a link between the increase in transcription of Orai3 and that of MRP-5. As for MDR1, MRP-5 seemsto be involved in resistance to cisplatin.

However, real-time quantitative RT-PCR does not provide any indication of the functionality of the proteins studied since an increase in the mRNA level does not necessarily reflect an increase in the number and functionality of the protein considered. Thus, an increase in the Orai3 transcript level does not mean that Orai3 will necessarily be in greater quantity and more functional. Only hypothesis (2) of the quench made it possible to make the link between an increase in the transcription of the Orai3 geneby pemetrexed and an increase in the number of functional membrane Orai3s.

The SYBRGreen binds to any double-stranded DNA, so the specificity of this technique is reduced. Another technique allows for greater specificity by using a Taqman probe coupled with the usual primers. If the primers bind to DNA other than that of interest, the synthesized double strand will notbe detected because the highly specific Taqman primer will not have bound to the DNA that is not of interest. This technique is, however, more expensive than the one using SYBR Green.The analysis of the melting curves (not shown here) allowed us in our case to ensure that only one PCR product was generated under the different conditions. To be certain that it was the PCR product of interest, itwould have been necessary to perform sequencing of the amplicon.

## CONCLUSION AND PERSPECTIVES

Our results show that pemetrexed increases the transcription rate of the Orai3 gene. We also suspected a possible transcriptional modulation by pemetrexed for other calcium channels. Wehighlighted a resistance to pemetrexed that seems to be independent of ABC transporters regulated by the Orai3 channel, mainly MDR1.

There are now molecular mechanisms that could be interesting to study in the phenomena of chemoresistance to pemetrexed. In particular, does the increase in the rate of Orai3 transcripts by pemetrexed have repercussions through a greater number of functional membrane Orai3 channels? A Western blot as well as patch-clamp can be undertaken to answer this question. The Western blot is performed with anti-Orai3 antibodies in order to assess the quantity of Orai3 channels with and withoutpemetrexed, after 72 hours of incubation. Since a greater number of membrane Orai3 channels is not necessarily linked to a greater functionality of these channels, the patch-clamp technique can be performed for the study at the functional level. It would be necessary to inhibit the maximum number of channels other than the one of interest, i.e. Orai3. For the remaining channels, the current generatedby Orai3 can be obtained using the RNA interference strategy directed against Orai3.

The current is recorded in SiCtrl and in SiOrai3 in both cases (with and without pemetrexed) then the difference in current, for each case, between SiOrai3 and SiCtrl gives the current generated by Orai3, allowing to see if the channel is more functional or not in the presence of pemetrexed. The absence of specific pharmacology for the Orai3 channel must indeed guide towards the use of the RNA interference strategy, requiring validation of all transfections before patch-clamp manipulations. Whether the increase in Orai3 transcript levels by pemetrexed is correlated with a greater number of membrane andfunctional Orai3 channels is an interesting issue since it directly conditions the effect of cisplatin when administered with pemetrexed, as is the case with Alimta co-treatment.and cisplatin. Indeed, if pemetrexed increases the rate of Orai3 transcripts but without increasing the number of functional Orai3 channels then for a given effective concentration of pemetrexed the co-administered cisplatin will be little subject to resistance phenomena, the quantity of functional membrane Orai3 not being amplified by pemetrexed. It should nevertheless be noted that this reasoning is valid if pemetrexed does not increase the number and functionality of ABC transporters responsible for the efflux of cisplatin, without passing through Orai3. Since in this case an increase in the number of functional Orai3 channels would not be necessary for this to result in greater resistance to cisplatin. On the contrary, if pemetrexed increases the number of functional membrane Orai3 channels then for this same concentration of pemetrexed, cisplatin will be more subject to resistance phenomena andits concentrations will have to be increased to have the same therapeutic effect, knowing that cisplatin also acts on the Orai3 channels and therefore increases the resistance of the cells to itself.

However, high concentrations of cisplatin are harmful to the patient. Another short-term research perspective would be to observe the phosphorylation rate of AKT inthe presence and absence of pemetrexed, since it has been suggested that pemetrexed can induce apoptosis via prolonged activation of AKT. A western blot can be used with an anti-p-AKT antibody.

Flow cytometry could be another complementary approach to determine the mode of action of pemetrexed. The MTT performed did not allow us to know whether pemetrexed decreased cell viability by promoting apoptosis, synchronization of cells in S phase or whether both phenomenacoexisted. Thus, annexin-V coupled to FITC could be used in flow cytometry to detect apoptosis if it occurs in the presence of pemetrexed. The mode of action of pemetrexed will allow us to know whether the resistance to pemetrexed is a resistance to cell mortality or synchronization or both.

The use of primary cultures for these studies also represents another research perspective. Our preliminary results suggest that Orai3 channels could ultimately constitute a new therapeutic target to improve the sensitivity of cancer cells to cisplatin and thus hope for a better life expectancy for patients with pulmonary adenocarcinoma. However, it would be necessary to carryout additional experiments to increase the sample size.

This internship allowed me to have a first approach to the professional world and public research in a laboratory attached to a faculty. Above all, this experience gave me a better knowledge of theexperimental techniques used and the different stages during manipulations. I was also able to discuss various hypotheses concerning my subject and propose several experimental protocols, which allowed me to appreciate the feasibility and the limit of these protocols, highlighting the difference between theoretical approaches and the experimental world. This allowed me to gain maturity as well as autonomy. In addition, this internship allowed me to discover how a team isorganized in a laboratory and more generally how the world of research works. All this helped to strengthen my idea of continuing in this field.

## Supporting information

https://sendeyo.com/show/26dcbc0924

## ABBREVIATIONS

| Abbreviation | Description |
| --- | --- |
| ABC | ATP Binding Cassette |
| MTT | 3-(4,5-dimethylthiazol-2-yl)-2,5-diphenyl tetrazolium bromide |
| RT-PCR | Reverse Transcription Polymerase Chain Reaction |
| MRP | Multidrug Resistance-associated Protein |
| LOL | Multidrug resistance |
| CBNPC | Non-Small Cell Lung Cancer |
| PI3K | Phosphoinositide 3 Kinase |
| MAPK | Mitogen-Activated Protein Kinases |
| KRAS | v-Ki-ras2 Kirsten Rat Sarcoma viral oncogene homolog |
| GRB2 | Growth factor Receptor-Bound protein 2 |
| SH | Src Homology |
| SOS | Son Of Sevenless |
| Ras-GEF | Ras-Guanine nucleotide Exchange Factors |
| Ras-GAP | Ras-GTPase Activating Proteins |
| Raf | Rapidly Accelerated Fibrosarcoma |
| MEK | Mitogen-Activated Protein Kinase Kinase |
| Erk | Extracellular signal-regulated kinases |
| HIF-1α | Hypoxia-Inducible Factor-1α |
| PMCA | Plasma Membrane Calcium ATPase |
| SERCA | Sarco/Endoplasmic Reticulum Calcium ATPase |
| CAMKK | Calcium/calmodulin-dependent protein Kinase Kinase |
| SOC | Store-Operated Channels |
| STEM | STromal Interaction Molecule |
| EMEM | Eagle's Minimal Essential Medium |
| HEPES | 4-(2-hydroxyethyl)-1-piperazineethanesulfonic acid |
| PBS | Phosphate Buffered Saline |
| EDTA | ethylenediaminetetraacetic acid |
| BAPTA | 1,2-bis(o-aminophenoxy)ethane-N,N,N,N-tetraacetic acid |
| DMSO | DiMethylSulfOxide |
| dNTP | DeoxyriboNucleotides TriPhosphate |
| cDNA | Complementary DNA |

