## Supplementary material for "Characterization and implication of the Orai3 channel and ABC type transporters in the phenomenon of chemoresistance to cisplatin and pemetrexed in lung cancer": https://sendeyo.com/show/26dcbc0924

Daoudi Redoane*

* University of Caen Normandy 14000 FRANCE

### SUMMARY

**ABBREVIATIONS.**

**ABSTRACT.**

**INTRODUCTION.	p.1**

**MATERIALS AND METHODS.	p.8**

**RESULTS AND DISCUSSION.	p.18**

**CONCLUSION AND PERSPECTIVES.	p.30**

**BIBLIOGRAPHY.**

### ABBREVIATIONS

Abbreviation

Description

ABC MTT RT-PCR MRP LOL CBNPC PI3K MAPK KRAS GRB2 SH SOS

Ras-GEF Ras-GAP

Raf MEK

Erk HIF-1*α* PMCA SERCA CAMKK SOC STEM EMEM HEPES PBS EDTA BAPTA DMSO dNTP cDNA

ATP Binding Cassette 3-(4,5-dimethylthiazol-2-yl)-2,5-diphenyl tetrazolium bromide

Reverse Transcription Polymerase Chain Reaction Multidrug Resistance-associated Protein

Multidrug resistance Non-Small Cell Lung Cancer

PhosphoInositide 3 Kinase Mitogen-Activated Protein Kinases

V- Ki-ras2 Kirsten Rat Sarcoma viral oncogene homolog Growth factor Receptor-Bound protein 2

Src Homology

Son Of Sevenless Ras-Guanine nucleotide Exchange Factors

Ras-GTPase Activating Proteins

Rapidly Accelerated Fibrosarcoma

Mitogen-Activated Protein Kinase Kinase Extracellular signal-regulated kinases Hypoxia-Inducible Factor-1*α*

Plasma Membrane Calcium ATPase Sarco/Endoplasmic Reticulum Calcium ATPase

Calcium/calmodulin-dependent protein Kinase Kinase

Store-Operated Channels STromal Interaction Molecule Eagle's Minimal Essential Medium

4-(2-hydroxyethyl)-1-piperazineethanesulfonic acid

Phosphate Buffered Saline

ethylenediaminetetraacetic acid

1,2-bis(o-aminophenoxy)ethane-N,N,N,N-tetraacetic acid

DiMethylSulfOxide

DeoxyriboNucleotides TriPhosphate

Complementary DNA

### ABSTRACT

**Orai3 channels have been associated with cell proliferation, survival and metastasis in several cancers. Previous studies have shown that Orai3 seems to be involved in the development of acquired chemo-resistance of cisplatin in non-small cell lung cancer via ABC transporters that transport certain chemotherapeutic agents, such as cisplatin, out of the cells. Based on these studies, we hypothesize that Orai3 may be involved in the development of acquired chemo-resistance of pemetrexed. By using MTT assay, we show that pemetrexed is efficient (IC50 =**0*,*4*µM***)and significantly decreases cell viability in**

After centrifugation and to remove the freezing medium, the supernatant obtained was removed and the A549 cells were resuspended in 1mL of culture medium.

The suspension was placed in a 25cm flask containing 5mL of culture medium. This flask was then kept in an incubator containing 5%*CO*2, an atmosphere saturated with humidity

and at 37˚C. After 24 hours of incubation, the cells adhere to the bottom of the flask and proliferate, forming a cell carpet.

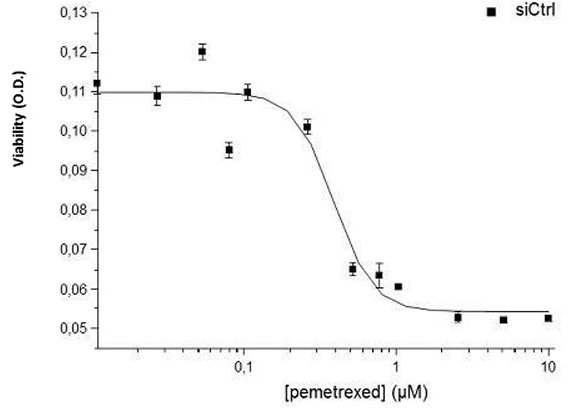

Figure 1 – Effect of pemetrexed on cell viability after 72 hours. (n=4)

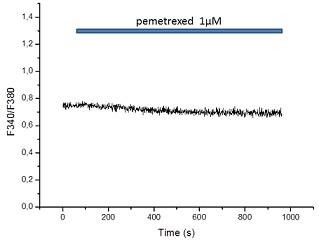

Figure 2 – Effect of pemetrexed on calcium channel activity.

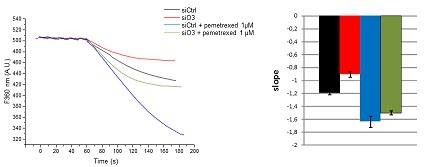

Figure 3 – Variations in membrane permeability to calcium with and without Orai3, without or with pemetrexed.

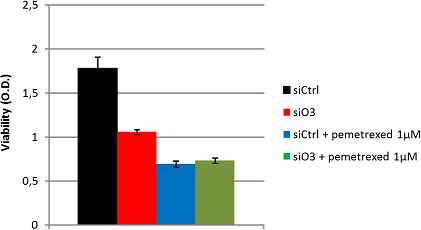

Figure 4 – Effect of pemetrexed on cell viability after 72 hours and in the absence of the Orai3 channel.

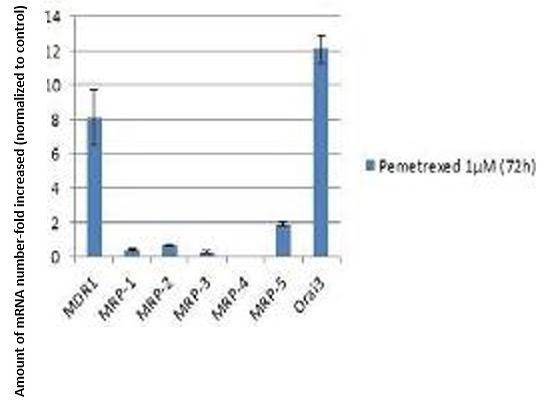

Figure 5 – Quantification of the mRNA level of ABC transporters and Orai3 in cells treated with pemetrexed (1µM, 72h)
